# LRLoop: Feedback loops as a design principle of cell-cell communication

**DOI:** 10.1101/2022.02.04.479174

**Authors:** Ying Xin, Pin Lyu, Junyao Jiang, Fengquan Zhou, Jie Wang, Seth Blackshaw, Jiang Qian

## Abstract

Genome-wide gene expression profiling, and single-cell RNA-seq in particular, allows the predictions of molecular mechanisms regulating cell-cell communication based on the expression of known ligand-receptor pairs. Currently available techniques for predicting ligand-receptor interactions are one-directional from sender to receiver cells. Here we describe LRLoop, a new method for analyzing cell-cell communication that is based on bi-directional ligand-receptor interactions, where two pairs of ligand-receptor interactions are identified that are responsive to each other, and thereby form a closed feedback loop. We assessed LRLoop using both bulk and single-cell datasets and found our method significantly reduces the false positive rate seen with existing methods. Finally, we applied LRLoop to the single-cell datasets obtained from retinal development and discovered many new bi-directional ligand-receptor interactions among individual cell types that potentially control proliferation, neurogenesis and/or cell fate specification.

## INTRODUCTION

Multicellular organisms rely on cell-cell communication to coordinate various biological processes and respond to environmental stimuli [1,2]. Decades of research have accumulated a massive amount of information on signaling pathways and ligand-receptor interactions. Recent sequencing technologies enable us to profile the gene expression at the single-cell level [3,4], and two major types of computational methods have been developed to mine single-cell RNA-Seq (scRNA-seq) data for biologically important cell-cell signaling interactions. The first type predicts the cell-cell communication based on the expression of ligands and their receptors [5–7]. If one cell type expresses a specific ligand and another cell type expresses its receptor(s), these two cell types are considered to be communicating with one another. The second class of methods uses entire transcriptomes to predict cell-cell communication [8,9]. By constructing the signaling networks and transcriptional regulatory networks, the downstream genes can be identified for each ligand. Gene sets that are differentially expressed in certain conditions can then be used to infer the upstream ligands.

These methods often generate large discrepancies in the predicted ligand-receptor interactions or patterns of cell-cell communication partly because they often have high false positive rates [10]. In this work, we propose a new method, ligand-receptor loop (LRLoop), which is based on the previous observation that signaling interactions between individual cell types are likely to form feedback loops. We assessed this in both bulk and single-cell RNA-seq datasets, and found that it outperformed other existing methods. Applying LRLoop to single-cell datasets obtained from developing mouse retina predicted many ligand-receptor interactions during retinal development.

## MATERIALS AND METHODS

### 1. Sources of ligand-receptor pairs, signaling and gene regulatory networks

- We combined literature-validated ligand-receptor pairs used in NicheNet [8] and connectomeDB2020[7]. We further filtered these ligand-receptor pairs using ligand and receptor annotation from CellTalkDB and NATMI database [7,11]. The ligand-receptor pairs were removed if the definition of the involved ligand or receptor was not supported by either of the two databases. After filtering, we retained 2,512 ligand-receptor pairs, involving 859 ligands and 726 receptors, for subsequent downstream analysis.
- To construct our default LRLoop network, we adopted the intracellular signaling network and gene regulatory network collected in NicheNet [8].

### 2. Construction of the LRLoop

The construction of LRLoop consists of three steps:

a. Construct the ligand/receptor-target regulatory potential matrices. With the collected ligand-receptor pairs, signaling and gene regulatory networks, we constructed the ligand/receptor-target regulatory potential matrices using NicheNet. For the calculation of these matrices, we adopted the functions construct_weighted_networks, apply_hub_corrections, construct_ligand_target_matrix in the R package nichenetr and a slightly modified version of the function construct_ligand_target_matrix (for the construction of the receptor-target matrix). The algorithm was based on the idea of propagation of the signal from a ligand/receptor to the downstream proteins mediating receptor signaling, transcriptional regulators targeted by these factors, and genes that are in turn regulated by these transcriptional regulators. Google’s Personalized PageRank algorithm was used to link the ligands/receptors to transcriptional regulators [8]. The resulting ligand/receptor-transcriptional regulator matrices were then multiplied to the transcriptional regulator-target matrix of the gene regulatory network to obtain the ligand/receptor and target gene relationships.
b. Identifying the target genes of each ligand/receptor. With the ligand/receptor-target regulatory potential matrices, we identified the set of target genes of each ligand/receptor which were defined as the ones with relatively higher regulatory potential scores among all the potential target genes of the ligand/receptor. For the construction of our default LRLoop network in the analysis, we used the function make_discrete_ligand_target_matrix of the nichenetr package with its default parameters (error_rate = 0.1, cutoff_method = “distribution”).
c. Finally, we identified all the [L1-R1]<->[L2-R2] pairs where L1 and L2 are among the target genes of each other for the feedback loops. Alternatively, we also identified the loops when L2 is a target gene of R1, and L1 is among the target genes of R2.

### 3. Assessment of LRLoop using bulk datasets

Each of the 111 ligand treatment datasets collected by Browaeys et al.[8] provides the treatment ligand, the expressed genes in the receiver cells and the differential expression information of these genes (including the logFC and q-value). For each of these datasets, based on our curated ligand-receptor network, we took the set of the ligand genes of all the expressed receptor genes in the receiver cells as the set of candidate ligands L1 and ranked these candidate ligands in the following ways:

- max{|logFC of R1|}: We scored each candidate ligand gene L1 by the value of max{|logFC(R1)|: R1 ∈ the set of cognate receptor genes of L1}. The ligands L1 are then ranked by these scores in descending order.
- CCCExplorer (L1 score): We replaced the ligand-receptor pairs and the gene regulatory network of CCCExplorer with the networks we used in the work so that the results obtained from different methods are comparable. As a required input of CCCExplorer, the built-in KEGG pathways remained the same. The candidate ligand genes L1 were then ranked based on a p-value score calculated by CCCExplorer in ascending order.
- NicheNet (L1 score): We took all the genes expressed in the receiver cells as background and those with q-value < 0.1 and the absolute value of logFC greater than or equal to 1 as the set of differentially expressed target genes to calculate the ligand activity scores of the candidate ligand genes L1 using nichenetr package. The ligand genes L1 were then ranked based on these scores in descending order.

For each method, we have also assessed the rank values for the treatment ligands using LRLoop filtering and random filtering. If a ligand did not form a feedback, we assigned the rank of the ligand to the last among all the candidate ligands.

### 4. Application of LRLoop to scRNA-seq data

We developed an interaction score (S_LR_) to quantify the strength of ligand-receptor interactions in a particular biological condition. The score considered the contribution from both L1-R1 and looped L2-R2 interactions between two cell types. Suppose L1-R1 is a candidate ligand-receptor interaction expressed from cell type A to cell type B (by default setting, detected in at least 10% of the cells in each cell type, respectively). Let w_L1R1_ be the interaction strength (e.g. gene expression, NicheNet score, or SingleCellSignalR score) of L1-R1 and w_L2R2_ the maximum of the interaction strength of its partners (L2-R2) expressed from cell type B to cell type A. If L1-R1 has no LRLoop partner, let w_L2R2_ = 0. The interaction score of the loop (L1-R1:L2-R2) was defined as:

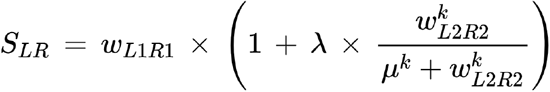

where 0 ≤ λ ≤1 can be understood as a weight for L2-R2 interaction, 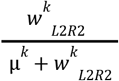 is a Hill function bounded between 0 and 1 that increases as the value of w_L2R2_ increases and the saturation curve becomes steeper as k > 0 gets larger. For the Mouse Cell Atlas data[12], the optimal parameters were obtained as λ = 0.9, μ = 0.7 and k = 4 when using the SingleCellSignalR score for the values of w_L1R1_ and w_L2R2_, and λ = 0.9, μ = 0.08 and k = 4 using NicheNet ligand activity score.

We adopted two methods to calculate the L-R interaction strength (i.e., w_L1R1_ and w_L2R2_):

- NicheNet (pearson): In each cell type, we took the genes that are detected in at least 10% of the cells as “expressed” genes and those whose average expression levels are above 75 (or 90, 95) percentile among all the expressed genes as “highly expressed”. We calculated the ligand activity score (the Pearson correlation coefficient version) for each expressed ligand using nichenetr package, taking all the expressed genes in the receiver cell type as background and the set of genes that are highly expressed in the receiver cell type as the target gene set of interest. An interaction L-R was counted if both L and R were expressed in the corresponding cell types and that the ligand activity score of L was above some threshold value.
- SingleCellSignalR: The interaction strength was defined as 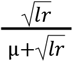 [5], where l is the average expression value of the ligand, r is the average expression value of the receptor, μ = *mean*(*C*) and C is the normalized read count matrix. We downloaded the high-quality batch-removed digital gene expression data (Mouse Cell Atlas) [12]. Tissues with only one cell type and cell types with less than 10 cells were removed for the analysis. For the remaining 36 tissues, we examined the proportions of between/within-tissue ligand-receptor interactions with or without consideration of L2-R2 contribution for each tissue pair at multiple cutoff score values.

### 5. Prediction of L-R interactions between retinal cells

We separately analyzed the ligand-receptor interactions for the three retinal development stages. We identified 5, 6, 6 retinal cell types in the early, intermediate and late stage, respectively. Based on the cell type annotations, we predicted the feedback LR loops and ranked them between each pair of these cell types using the LRLoop package.

Before running LRLoop, we first cleaned the ligand-receptor pairs according to their annotation in the Human protein atlas [13] and Uniprot [14]. We removed the ligand-receptor pairs if one of them has been annotated as “intracellular proteins”in both datasets.

Next, we detected feedback loops between each pair of cell-types following the standard LRLoop analysis pipeline. Briefly, to calculate feedback loops between cell-type A and cell-type B, we first cleaned the LR networks based on the ligand and expression (at least 2.5% cells expressed) in cell-type A, and the receptors expression (at least 2.5% cells expressed) in cell-type B. We then cleaned the signaling networks based on the gene expression (at least 2.5% cells expressed) in cell-type B. With the cleaned LR network, cleaned signaling network and the gene regulatory network, we constructed 2 matrices: AtoB-ligand-target matrix and

AtoB-receptor-target matrix. Conversely, with the same cleaning method, we constructed another 2 matrices: BtoA-ligand-target matrix and BtoA-receptor-target matrix. With the 4 matrices, we detected the feedback loops using the function: “PrepareBasics” with min_pct = 0.025.

Finally, for each pair of cell-types, we calculated the S scores for all detected ligand-receptor pairs. To quantify the global interactions between each cell-type pair, we calculate the aggregated *S_c_* score, which was defined as:

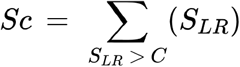

where C is the cutoff for S_LR_. To find the more specific LR pairs, we further filtered the ligand-receptor pairs with the following criteria: 1) we removed LR pairs if they are present at least 70% cell-type pairs. 2) We removed the LR pair if their maximum difference of S_LR_ score is < 0.5 across all cell-type pairs.

## RESULTS

### Closed feedback loop as a design principle for cell-cell communication

Here, we propose a new method, ligand-receptor loop (LRLoop), which identifies feedback loop signaling interactions between individual cell types. Specifically, for the two-way communication to occur, cell type A secretes ligand L1, which interacts with receptor R1 in cell type B. In turn, cell type B responds to the signal from cell type A by secreting ligand L2 that interacts with receptor R2 in cell type A. To make the L1-R1 and L2-R2 interactions responsive to each other, the ligands (L1 and L2) are required to be encoded by the genes that are regulated by receptors in the corresponding cells (Figure 1A). We indeed observed instances of this type of design principle in controlling cell-cell communication. For example, during injury-induced Muller glia reprogramming in zebrafish [15], microglial cells secrete ligand Tnf, which binds to receptor Tnfrsf1a, which in turn is expressed in Muller glial cells. At the same time, the ligand Anxa2 is secreted by Muller glial cells, and its receptor Tlr2 is expressed in microglial cells. Furthermore, based on known signaling and regulatory networks, Tnf and Anxa2 are predicted to be downstream genes of Tlr2 in microglia and Tnfrsf1a in Muller glia, respectively. These four genes are found to be co-expressed in the time course; they are all upregulated at 3 hours after retinal injury, and gradually downregulated afterward (Figure 1B) [15]. The Tnf-Tnfrsf1a mediated microglia-Muller glia communication was confirmed in zebrafish retina [16]. Our analysis suggests that the feedback loops might be one type of design principle regulating cell-cell communication.

**Figure 1.**
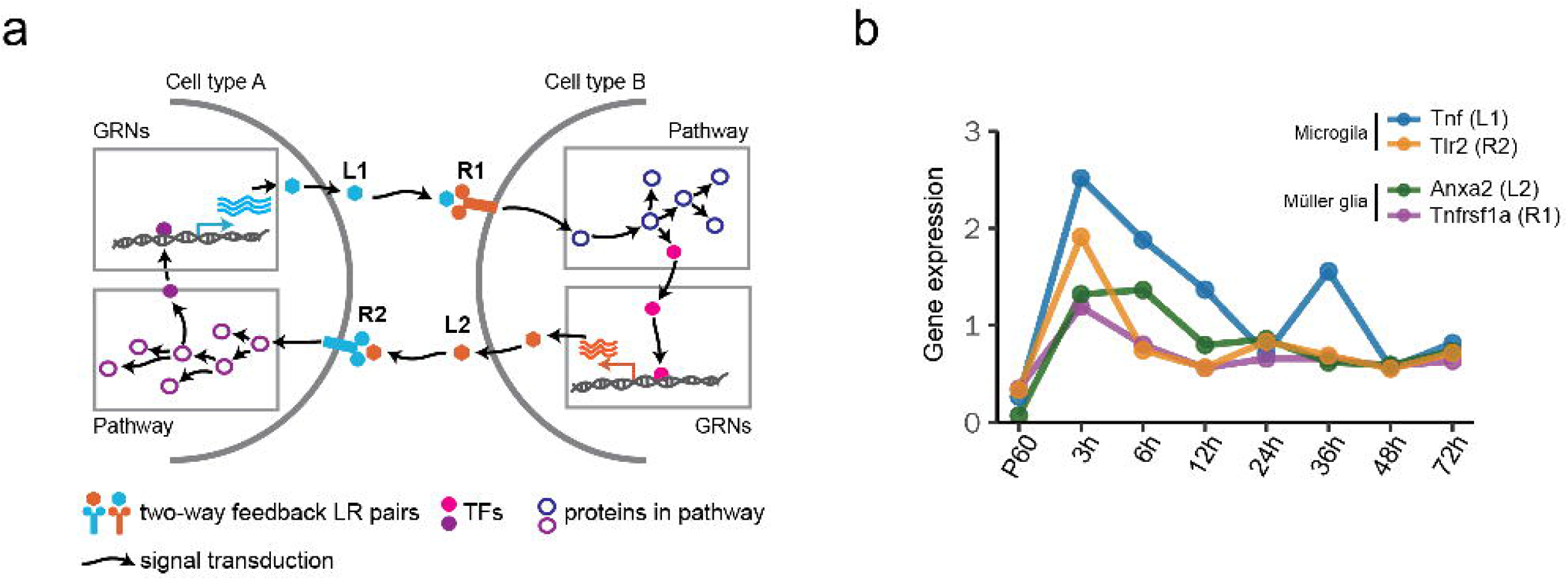
LRLoop algorithm and its assessment. **a.** Schematic of the LRLoop concept. Ligand 1 (L1) interacts with receptor 1 (R1), while ligand 2 (L2) interacts with receptor 2 (R2). Furthermore, L1 is the downstream target of R2 through signaling pathways and gene regulatory networks (GRNs). Similarly, L2 is the downstream target of R1. **b.** An example feedback loop between Muller glia and microglia.

We next investigated the prevalence of the feedback loops based on known ligand-receptor pairs, signaling pathways, and gene regulatory networks. In total, we identified 2,512 experimentally supported ligand-receptor pairs based on literature. Among all 3,156,328 possible loops involving two known ligand-receptor pairs (L1-R1 and L2-R2), we found that 0.55% (17,413/3,156,328) of them form feedback loops (see Methods, Figure 2A). These pairs involve 656 ligands and 647 receptors. The majority of known ligand-receptor pairs (77%, 1935/2512) were found in at least one signaling feedback loop. Most ligand-receptor interactions form the feedback loops with a few other ligand-receptor interactions, while a few of them could partner with many interactions (Figure 2B). The top ligand-receptor interaction identified is VCAN-EGFR, which pairs with 600 other ligand-receptor interactions to form feedback loops, probably in various biological contexts.

**Figure 2.**
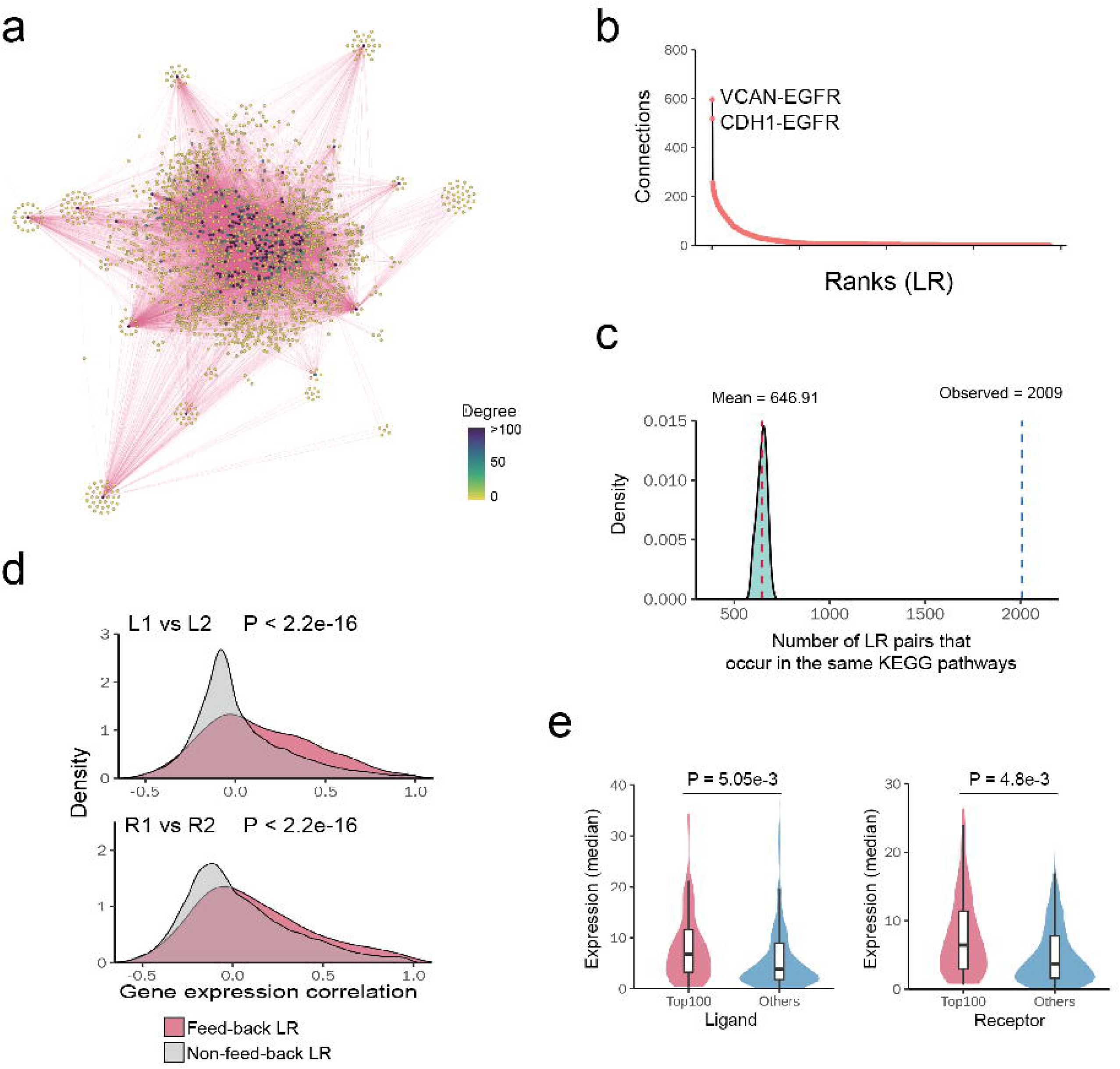
Properties of identified ligand-receptor loops. **a.** Networks of feedback ligand-receptor pairs. Each node represents a ligand-receptor pair. Two nodes are connected if they form a feedback loop.**b.** Ranked ligand-receptor pairs based on their connection degree in the feedback network (high to low). X axis indicate their ranks. Y axis indicate their number of connections. **c.** Number of the feedback loops whose four members (L1, R1, L2 and R2) are in the same KEGG pathways. The distribution shows the same numbers if we randomly select two ligand-receptor pairs. **d.** The Pearson correlation coefficients of connected L1-L2/R1-R2 are significantly higher than un-connected L1-L2/R1-R2. P values were calculated using KS-test. **e.** The median expression level of hub ligands/receptors are significantly higher than lowly connected ligands/receptors. P values were calculated using t-test.

Several lines of evidence suggest that the feedback loops identified using this approach are biologically relevant. First, we assessed whether the LR loops are more likely to be in the same biological pathways. In fact, members of 2,009 feedback loops (L1, L2, R1, and R2) are found in the same KEGG pathways. For example, LAMA2-ITGB1 and PDGFB-PDGFRA interactions are found to form a feedback loop, and both pairs of the proteins are part of the focal adhesion signaling pathway [17]. In contrast, randomly selected pairs of ligand-receptor interactions are much less likely to be in the same KEGG signaling pathways (Figure 2C). Second, we examined whether the genes encoding the feedback loop members tend to be co-expressed. We analyzed the gene expression profiles across human tissues and calculated the correlation coefficients of gene expression[18]. Specifically, we calculated three correlation coefficients (L1-L2, R1-R2) and found that they were higher than those obtained from random pairs of ligand-receptor interactions (p<2.2e-16) (Figure 2D). Third, we examined whether ligand-receptor hubs have higher expression levels than non-hub genes, since these hubs formed feedback loops with many other ligand-receptor interactions and were more likely to be activated in multiple cell types or physiological conditions. We calculated the average gene expression of ligand-receptor pairs across human tissues, and found that the genes in the hubs were higher expressed than non-hub genes (p = 5.05e-3, p=4.8e-3) (Figure 2E). In summary, these results suggest that the feedback loops might be one widespread design principle regulating cell-cell communication.

### Assessment of LRLoop using bulk datasets

We next assessed whether the requirement of the existence of feedback loops could improve the prediction by incorporating the identified feedback loops to different scoring methods (e.g., gene expression changes or ligand activity score from NicheNet [8]). We first applied our approach to a transcriptome dataset compiled by Browaeys et al. in which the gene expression from bulk samples was profiled before and after the cells were stimulated by one or two ligands [8]. The dataset includes the transcriptomes from 111 treatments with a total 121 ligands. Note that the data only contain the expression profiles from the receiver cells, which include the differential expression data for R1 and L2 (Figure 3A). We examined the rank of the treatment ligands (i.e., the expected ligands) among all ligands in the system using different scoring methods. When we ranked the ligands based on the gene expression changes of their interacting receptors (R1) before and after the treatment, the median of the rank for the expected ligands was 179, suggesting that on average 178 other ligands rank better than the expected ligand. If we required both R1 and L2 to be present and ranked the ligands based on the expression changes of R1, the rank of the expected ligands was improved, with a median value of 114. In contrast, if we used randomly created feedback loops as a filter, the median rank of expected ligands was 383 because many expected ligands (59 on average) did not form a feedback in the random dataset and were ranked at the bottom (Figure 3A). Similar results were observed when we ranked the ligands based on the ligand activity scores using CCCExplorer[9] and NicheNet[8]. When we used the LRLoop as a filter, our method improved the rank of the expected ligands. In summary, our results indicated that the feedback loops could be incorporated into different scoring methods and substantially reduce the false positive rate.

**Figure 3.**
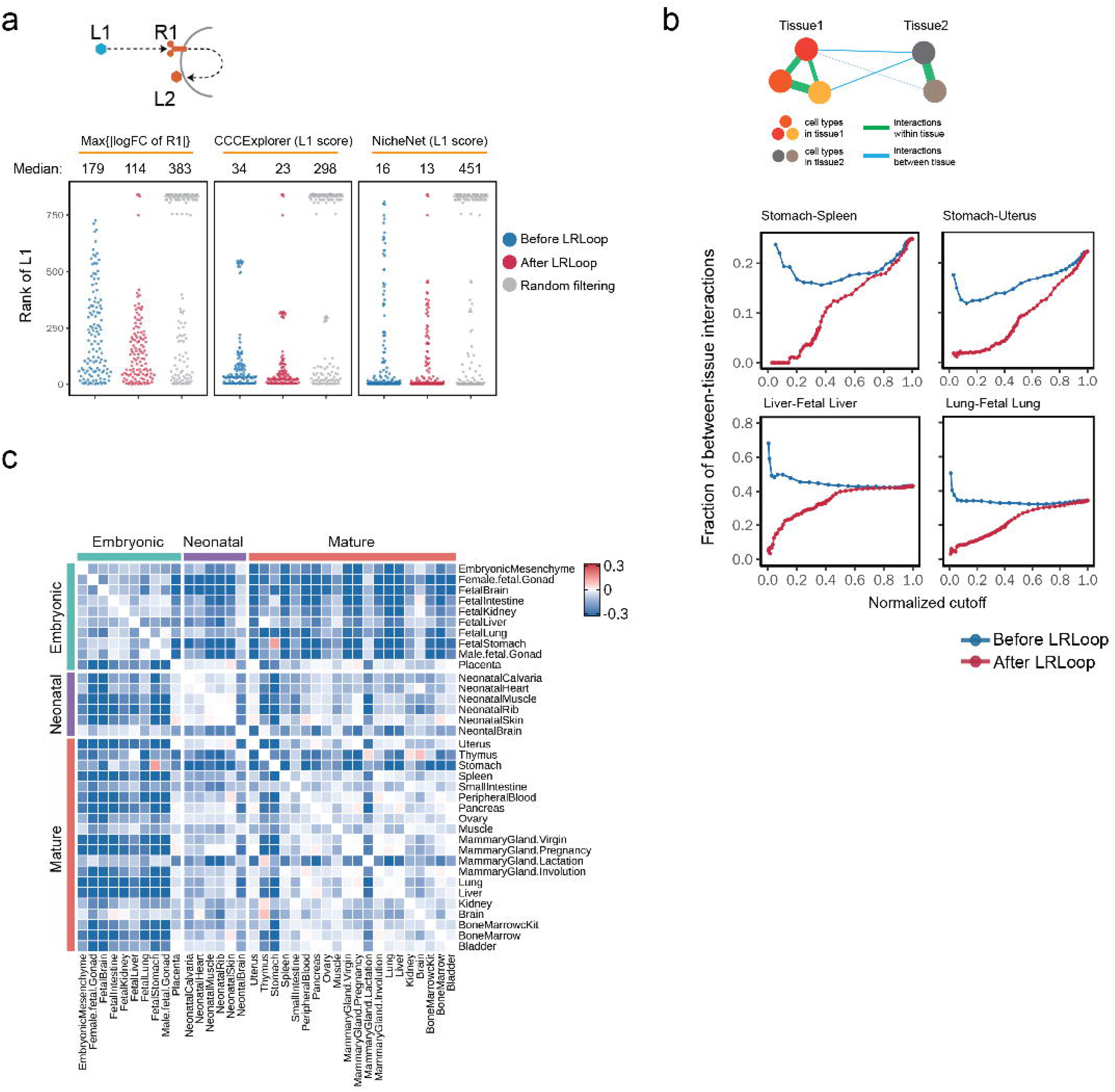
Assessment of LRLoop using bulk and single-cell datasets. **a**. Assessment using bulk expression datasets. Each dot represents an expected ligand. **b.** Assessment using single-cell datasets. We used the between-tissue interactions as the indicator for false positive predictions. In the x-axis are the fractions of within-tissue interactions that remain in all expressed within-tissue interactions above the cutoff score values. The y-axis displays the fractions of between-tissue interactions in all interactions that remain at the cutoff score values. **c**. Heatmap showing the reduction of between-tissue interactions, calculated as the ratio of the area under the “After LRLoop” curve to the area under the “Before LRLoop” curve as in b, and then minus one, among 630 tissue pairs.

### Assessment of LRLoop using single-cell datasets

We then systematically assessed our approach using scRNA-seq datasets. One challenge is that a comprehensive gold standard does not exist to evaluate the performance of the predicted patterns of cell-cell communication. To benchmark the prediction performance, we reasoned that the cells that were either spatially or temporally separated are less likely to signal to one another (Figure 3B). For example, cells in the spleen and stomach are less likely to interact with each other because these cells are spatially separated. Likewise, cells in fetal lung and mature lung are also less likely to interact because they are temporally separated. For a pair of tissues, we calculated the numbers of predicted ligand-receptor interactions within tissues and between the two tissues, and used the fraction of between-tissue interactions among all interactions as an indicator of the false positive rate.

To quantitatively include the contribution from both L1-R1 and L2-R2 interactions, we developed an empirical interaction score (S_LR_) to quantify the strength of ligand-receptor interactions (see Methods). We then applied the method to scRNA-seq data from mouse tissues at embryonic, neonatal and mature stages [12]. We found that incorporating the feedback loops could reduce the rate of predicted between-tissue interactions. For instance, the stomach includes stomach epithelial cell, gastric mucosa cell and endocrine cell types, while the spleen includes neutrophil, monocyte, dendritic cell, macrophage and NK cell types. We then calculated the interaction score based on the SingleCellSignalR[5] score. When we used different S_LR_ values as cutoffs, we found that the fraction of between-tissue interactions ranged from 0 to 0.25. If we do not consider the contribution from L2-R2 interactions (i.e., W_L2R2_= 0, see Methods), the corresponding range was from 0.15 to 0.25 (Figure 3B). The fraction of between-tissue interactions was substantially reduced if we considered the contribution from L2-R2 interactions. Similarly, we also observed the reduction of between-tissue interactions for temporally separated tissues. For example, the interactions between fetal liver and adult liver as well as fetal lung and adult lung, the fraction of between-tissue interactions was significantly reduced after we included the looped L2-R2 (Figure 3B). Overall, we observed 96.7% of 630 pairs of 36 tissues showed a reduction in the fraction of between-tissue interactions with an optimized parameter set (Figure 3C). Similar results were observed when we applied the feedback loops to the NicheNet scoring method (Supplementary Figs. 1a-c). In summary, our results suggest that the LRLoop approach can reduce the false positive rate obtained from analysis of single-cell datasets.

### Cell-cell communication during retinal development

Previous studies have highlighted the importance of cell-cell signaling during retinal development, with extrinsic factors controlling progenitor proliferation, neurogenesis, neuronal differentiation, and synapse formation [19,20]. We applied our method to retinal single-cell RNA-seq datasets and predicted cell-cell communication between individual retinal cell types during retinal development[21]. The scRNA-seq datasets are grouped into 3 different stages of neurogenesis: early (E14-E16), intermediate (E18-P2) (Supplementary Figure 2A) and late (P5-P8) (Supplementary Figure 2B). During early stages of neurogenesis, five distinct cell types were identified, including retinal progenitor cells (RPCs), amacrine cells/horizontal cells (AC/HC), retinal ganglion cells (RGCs), cone photoreceptors, and neurogenic cells (Figure 4A). We identified the ligand-receptor feedback loops between each pair of cell types. A cell-cell communication map was obtained by summarizing all ligand-receptor interactions (Figure 4B). Strong interactions were observed between RPCs and other cell types. Among these, 38 ligand-receptor interactions were identified for RPC-RPC signaling (Supplementary Figure 2C), which included a number of previously-identified signaling interactions. For example, fibroblast growth factor (FGF) signaling is known to play a key role in retinal progenitor proliferation and neurogenesis [22]. Consistent with previous knowledge, Fgf9-Fgfr1, Fgf15-Fgfr1, Fgf3-Fgfr1 interaction was found in retinal progenitor cells. Similarly, 11 known ligand-receptor interactions were identified between RPC and neurogenic cells, including multiple Notch-Delta pathway members (Supplementary Figure 2D) [23]. Interactions identified between RGC and neurogenic progenitors included Sst-Sstr2, which is known to play a role in photoreceptor generation and regulation of retinal neurogenesis [24] (Figure 4C, D).

**Figure 4.**
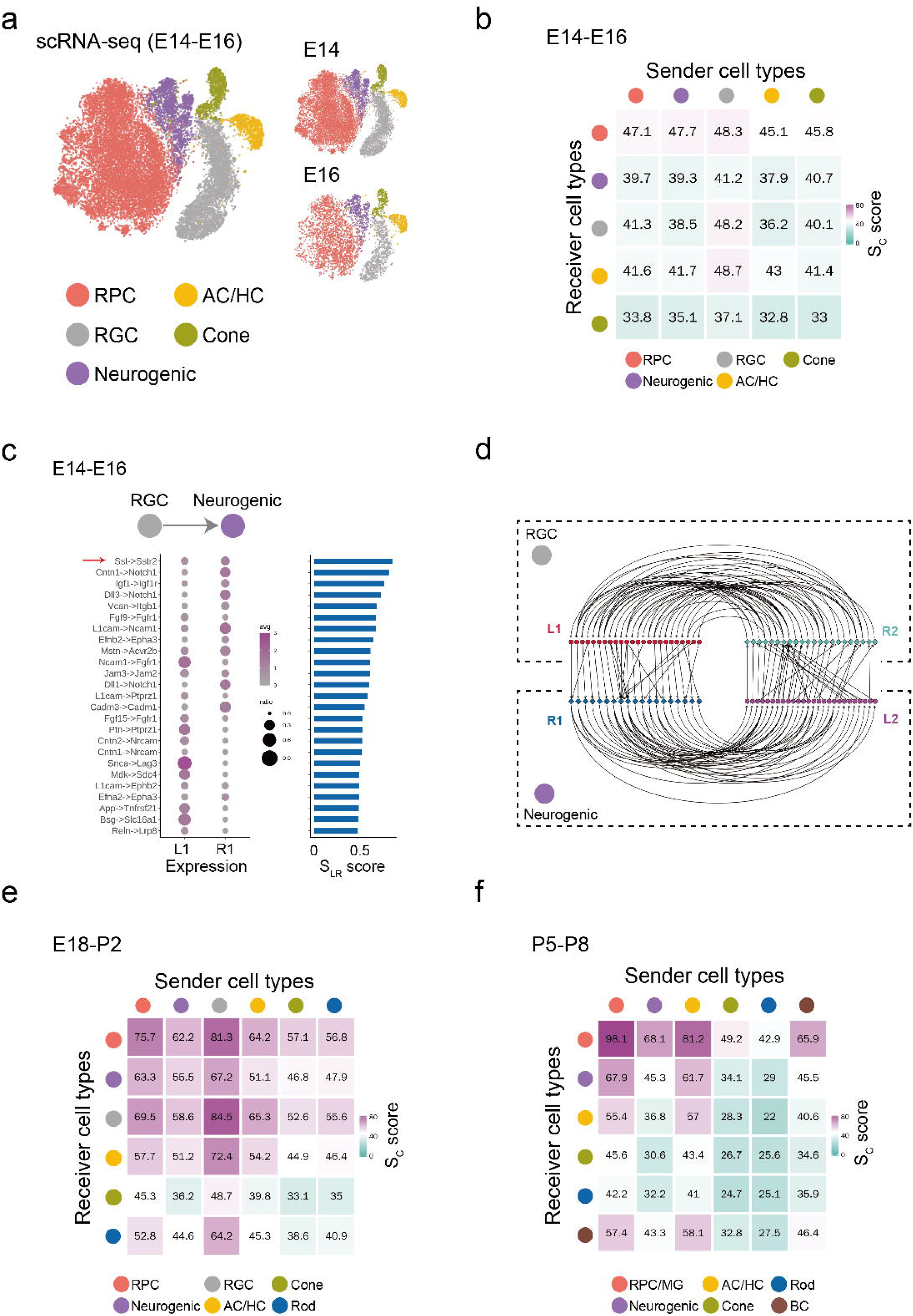
Application of the LRLoop to retinal development. **a.** Five major cell types identified using scRNA-seq dataset during the early retinal developmental stages (E14 and E16). **b.** Sc score between the five cell types calculated using LRLoop during early stages. **c.** The unique ligand-receptor interactions predicted between retinal ganglion cells (RGCs) and neurogenic cells during early stages. The red arrow indicates the known interactions. **d.** Detailed feedback loops for interactions between retinal ganglion cells (RGCs) and neurogenic cells during early stages of retinal neurogenesis. **e.** Sc score between the six cell types calculated by LRLoop during intermediate stages. **f.** Sc score between the six cell types calculated by LRLoop during late stages.

The patterns of cell-cell communication changed substantially over the course of retinal development. While during early neurogenesis, signaling between RPCs and other cell types was most prominent, during intermediate stages of neurogenesis, the number of all interactions increased substantially. In particular, many interactions were observed between RGCs and AC/HC, as well as between RPCs and other cell types (Figure 4E). During late neurogenesis, signaling between AC/HC and BC was prominent, as were reciprocal signaling between AC/HC and RPC/MG (Figure 4F, Supplementary Table 1). Both the timing and pattern of cell-cell signaling at later developmental stages coincides with the establishment of synaptic connectivity [20]. As expected, signaling loops between GABAergic AC/HC and glutamatergic BC and RGCs include both multiple Neuroligin-Neurexin and Cadm gene family members, which mediate formation of GABAergic and glutamatergic synapses, respectively [25,26] (Supplementary Table 1).

## DISCUSSION

Cell-cell communication plays an essential role in many biological processes in multicellular organisms. While many computational methods were developed for the prediction of ligand-receptor interactions, a recent survey of this field highlighted two major unsolved challenges [10]. First, a large discrepancy exists between the results obtained from different prediction methods. Second, we do not have a good gold standard dataset to assess the accuracy of these predictions. In this study, we developed a new computational method, LRLoop, to predict bi-directional ligand-receptor interactions. Compared to previous existing methods, the unique aspect of our method is that we require the presence of two ligand-receptor pairs that form a feedback signaling loop between two individual cell types. Furthermore, the expression regulation of the two ligands is dependent on the activity of the two receptors. This design principle of cell-cell communications ensures a robust bi-directional communication between cell types.

This feedback loop was identified in studies of cell-cell communication [27–29]. However, several key points are different in our analysis. First, in previous work, the two ligand-receptor interactions that form the feedback loop were considered to be independent of one another. The two ligand-receptor interactions were identified independently, using one-directional algorithms. In contrast, the two ligand-receptor interactions identified by LRLoop were responsive to one another, and are interconnected through previously validated signaling networks and regulatory networks. Second, in previous work, the feedback loops were found only after the ligand-receptor interactions were identified, while the feedback loops were used as a prerequisite for the prediction of ligand-receptor interactions in this study.

We expect that the closed feedback loop is a common design principle regulating cell-cell communication. Other design principles, potentially involving more than two cell types, are possible. The lack of a full set of biologically validated receptor-ligand relationships could be one possible reason for the false negatives observed in the bulk dataset. Furthermore, if additional relevant information is included, such as a more detailed analysis of the spatial relationships between cell types, it should be possible to more accurately predict patterns of cell-cell communication.

## CODE AVAILABILITY

The LRLoop R package is available at https://github.com/Pinlyu3/LRLoop

## ACKNOWLEDGEMENTS

We thank Drs. Mandeep Singh and Jeff Mumm for discussions. The work was partially supported by NIH grants (5R01EY029548 and 5P30EY001765 to J.Q.).

## Figure legends

**Supplementary Figure 1. Heatmaps showing the reduction of between-tissue interactions when applying LRLoop to NicheNet ligand activity scores for tissue pairs of the mouse cell atlas scRNA-seq data [21]**.

Both L and R are detected in at least 10% of the cells in the corresponding cell types. In the calculation of the ligand activity scores of the ligands, we take all the genes detected in more than 10% of the receiver cells as background and take highly expressed genes (genes whose average expression levels are above 75 (**a**), 90 (**b**) or 95 (**c**) percentile among the background genes) in the receiver cell type as the target gene set of interest. Reduction values are then calculated in the same way as in Figure 3C.

**Supplementary Figure 2. Application of LRLoop methods to the retinal development datasets. a**. TSNE plot showing the six major cell types in the single-cell RNA-seq datasets during intermediate stages of retinal neurogenesis (embryonic day 18-postnatal day 2). **b**. TSNE plot showing the six major cell types in the single-cell RNA-seq datasets during late stages of retinal neurogenesis (postnatal day 5-8). **c**. The unique ligand-receptor interactions predicted between retinal progenitor cells (RPC) during early stages. The red arrows indicate the known interactions. **d**. The unique ligand-receptor interactions predicted between neurogenic cells and RPCs during early stages.

