## Supplementary figures and images for "LRLoop: Feedback loops as a design principle of cell-cell communication"

### Supplementary Figures.docx

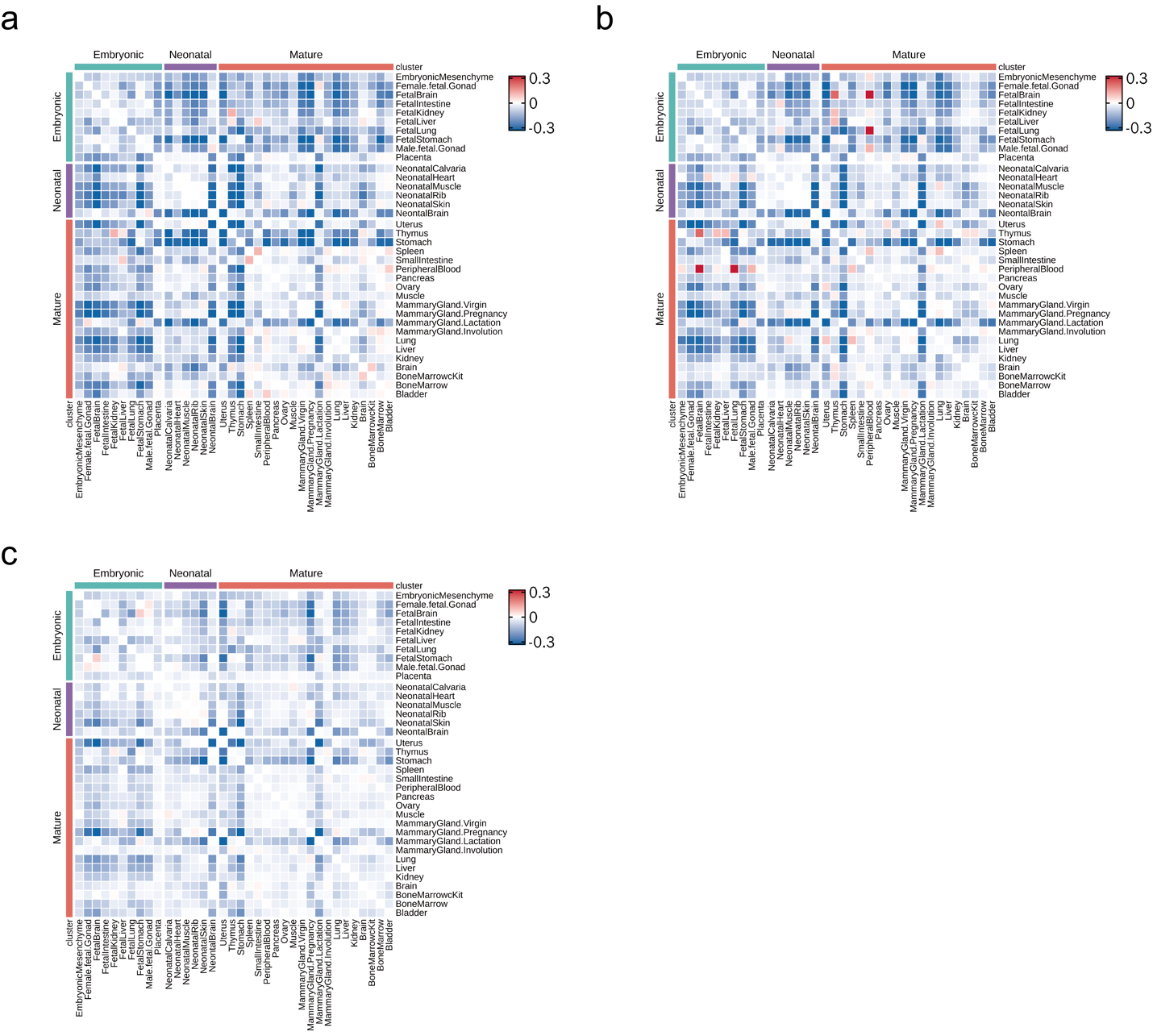

Supplementary Figure 1

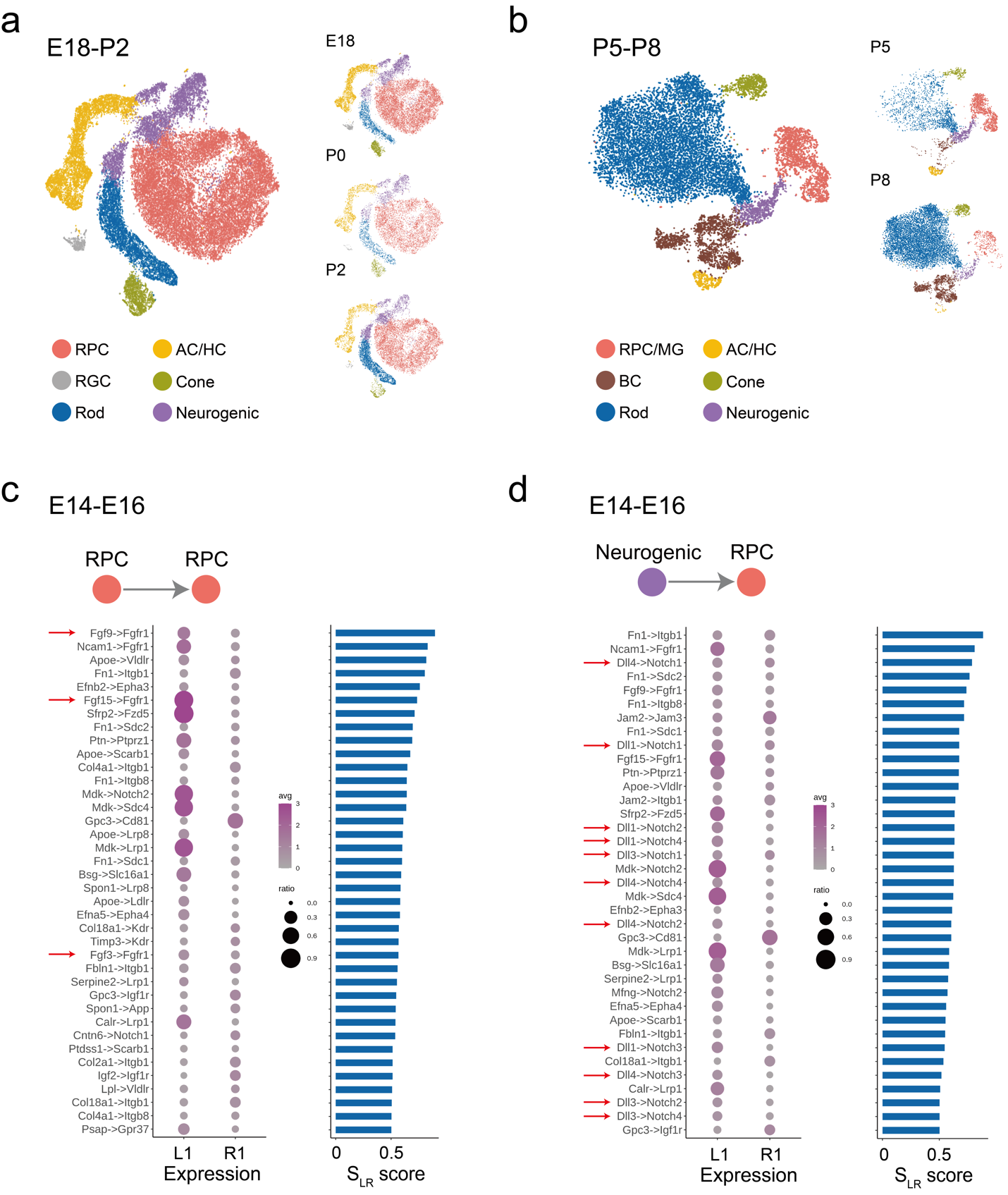

Supplementary Figure 2
